# Towards an Objective Evaluation of EEG/MEG Source Estimation Methods: The Linear Tool Kit

**DOI:** 10.1101/672956

**Authors:** Olaf Hauk, Matti Stenroos, Matthias Treder

## Abstract

The question “What is the spatial resolution of EEG/MEG?” can only be answered with many ifs and buts, as the answer depends on a large number of parameters. Here, we describe a framework for resolution analysis of EEG/MEG source estimation, focusing on linear methods. The spatial resolution of linear methods can be evaluated using the resolution matrix, which contains the point-spread and cross-talk functions (PSFs and CTFs), respectively. Both of them have to be taken into account for a full characterization of spatial resolution. They can be used to compute a range of quantitative resolution metrics, which should cover at the last three aspects of those functions: localization accuracy, spatial extent, and relative amplitude. Here, we first provide a tutorial-style introduction to resolution analysis of linear source estimation methods. We then apply these concepts to evaluate the benefit of combining EEG and MEG measurements, and to compare weighted and normalized L2-minimum-norm estimation and spatial filters. We confirm and extend previous results, showing that adding EEG to MEG improves spatial resolution, and that different methods offer different compromises among different resolution criteria. We hope that our approach will help EEG/MEG researchers in the interpretation of source estimation results, the design of new experiments, and the development of new MEG systems.

## Introduction

Electro- and magnetoencephalography (EEG and MEG) record electrical brain activity with high temporal resolution (in the millisecond range). Thus, their signals are directly related to the computational brain processes of interest to neurophysiologists and cognitive scientists, because they can track and distinguish fast perceptual and cognitive processes as well as brain oscillations. However, it is well known that these strengths of EEG and MEG are accompanied by a considerable weakness with respect to their spatial resolution. The question “What is the spatial resolution of EEG/MEG?” can only be answered with many ifs and buts, as the answer depends on a large number of parameters such as the measurement configuration, source estimation method and its assumptions, head model, source positions, signal-to-noise ratio, etc. Here, we describe a comprehensive and powerful framework to answer this question for linear source estimation methods.

Even if it were possible to perfectly record full three-dimensional electric and magnetic field distributions around the head, it would be impossible to unambiguously determine the three-dimensional source current distribution inside the brain that gave rise to those measurements (Sarvas, 1987); this inverse problem is ill-posed. Thus, unlike for example for fMRI, spatial resolution is not just limited by technology, but also by the physical principles governing signal generation.

A number of methods have been developed to find possible solutions to this inverse problem (Baillet, Mosher, & Leahy, 2001; Michel et al., 2004; Wipf & Nagarajan, 2009). All those methods are at their best when their assumptions are exactly fulfilled. For example, in cases where it is certain that the signal is generated by one dominant focal current source, then the source location, orientation and amplitude can be estimated with reasonable accuracy using a single equivalent dipole fit (Lütkenhöner, 1998b; Scherg & Berg, 1991). Unfortunately, in many EEG/MEG experiments, especially those with low signal-to-noise ratios (SNRs) or involving complex brain processes, it is impossible to specify prior assumptions on source activity in a precise mathematical manner. In those cases, it is common to employ distributed source solutions, which attempt to explain the measured signal as source-current distributions with the whole brain volume or across the cortical surface.

It is generally accepted that distributed-source solutions do not accurately reconstruct the true source distribution, but rather produce a “blurred” version of it. The degree of this blurring, i.e. the spatial resolution of these methods, and how it is affected by parameters such as sensor configuration, source depth, or the method itself, is still largely unknown. Commercial and freeware software packages for EEG/MEG analysis offer a large selection of source estimation methods, but rarely the tools to objectively evaluate them (Baillet, Friston, & Oostenveld, 2011; Dalal et al., 2011; Delorme & Makeig, 2004; Gramfort et al., 2013; Litvak et al., 2011; Oostenveld, Fries, Maris, & Schoffelen, 2011; Tadel, Baillet, Mosher, Pantazis, & Leahy, 2011).

Spatial resolution is a key concept for EEG/MEG source estimation. The main purpose of many EEG/MEG studies is to make inferences about the spatio-temporal brain dynamics of perceptual and cognitive processes; to achieve this, statistical analysis of source estimates is essential. Source estimation is also often used as an intermediate processing step in order to apply time series analysis, such as connectivity metrics, to the source time courses. In all of these cases, having a realistic idea about the spatial resolution of the source estimates is key for the interpretation of the results.

The evaluation of spatial resolution is more complicated for EEG/MEG than it is for fMRI, as it depends on more parameters such as the source estimation method (e.g. the type of depth weighting or noise normalisation), source location (e.g. depth), measurement configuration (number and location of magnetometers, gradiometers, EEG electrodes), the head model (spherically symmetric, realistic single-shell, three-shell, high-detail), as well as the SNR and noise covariance of the data (which may vary over time in any given data set). Researchers therefore need tools for a comprehensive evaluation of the spatial resolution of their experimental setup at hand.

The most popular EEG/MEG source estimation methods are arguable of L2-minimum-norm type (e.g. classical (optionally weighted) L2-minimum-norm estimation, L2-MNE; dynamic statistical parametric mapping, dSPM; standardized and exact low resolution electromagnetic tomography, s/eLORETA), dipole models, and linearly constrained beamformers. These methods have in common that they result in linear transformation of the data (with some caveats discussed later). They are applied to the data by means of vector or matrix multiplication to produce an estimate of the strength of a source at a particular location (“spatial filter”) or of the whole source distribution, respectively.

Linear transformations have some convenient properties that can be exploited for the evaluation of their spatial resolution (Backus & Gilbert, 1968; Dale & Sereno, 1993; Hauk, Stenroos, & Treder, 2019; Menke, 1989). Most importantly, the superposition principle holds, i.e. the source estimate of a signal produce by multiple sources is the sum of the source estimates produced by each individual source. Thus, knowing the source estimates for all possible point sources (i.e. their point-spread functions, PSFs) enables us, in principle, to construct the source estimate for any arbitrary configuration of sources. In order to see why this is useful, imagine a case where we are interested in two brain locations, and we would like to know whether activity originating from these two locations would produce peaks around the correct locations, and whether the peaks from the two locations could be distinguished from each other. We could test this by computing PSFs at each location, and evaluate their overlap. An important property of this approach is that is does not rely on the assumptions originally employed in the design of the source estimation method. No matter where we found a linear transformation, PSFs will demonstrate its performance. This framework is therefore well-suited to provide an objective and unbiased evaluation of spatial resolution for linear methods. Some important caveats with respect to the applicability of the superposition principle to (non-linear) intensity distributions and beamformers will be discussed below (see also Hauk et al., 2019).

Another key concept for the evaluation of source estimation methods is that of cross-talk: When we estimate the activity one source at a particular location, how much is it affected by activity from other locations? This is described by cross-talk functions (CTFs), which for a particular source describe how all other possible sources with unit strength would affect the source estimate for this source of interest. This concept is especially relevant for the application of spatial filters (or “software lenses”) that are supposed to filter out activity from a particular brain location, while suppressing activity from other locations (Grave de Peralta Menendez, Gonzalez Andino, Hauk, Spinelli, & Michel, 1997; Hauk et al., 2019; Van Veen, van Drongelen, Yuchtman, & Suzuki, 1997). While the concepts of PSFs and CTFs are related (they describe “leakage” in source space), there are important differences that will be discussed below.

It is of course cumbersome to look at all possible PSFs and CTFs individually. Thus, it is necessary to define resolution metrics that summarize important features of these distributions, which can then be visualized as whole-brain distributions, and compared across methods and across the parameter space. This has been done previously to evaluate the benefit of combining EEG and MEG measurements (Molins, Stufflebeam, Brown, & Hamalainen, 2008) to compare noise-normalized MNE methods (Hauk, Wakeman, & Henson, 2011), to compare different head models (Stenroos & Hauk, 2013), and to compare different MEG sensor types (Iivanainen, Stenroos, & Parkkonen, 2017).

Three aspects of spatial resolution are particularly important for most applications of EEG/MEG source estimation. First, the peaks of PSFs and CTFs should be close to the true source location. This has often been evaluated by means of the peak localization error, i.e. the Euclidean distance between the estimated peak and the true source location. Second, in order to be able to evaluate to which degree two distributions overlap with each other, we need a measure of spatial extent. This has for example been done using the standard deviation around the peak of the distribution, also called spatial dispersion. Third, even if two distributions are well-localized and largely non-overlapping, they might substantially differ in amplitude. Thus, if two sources are active simultaneously, one may completely overshadow the other. Thus, we need a measure for the relative amplitude of these distributions.

A number of previous studies have evaluated linear source estimation methods using different metrics (e.g. Fuchs, Wagner, Kohler, & Wischmann, 1999; Grave de Peralta-Menendez & Gonzalez-Andino, 1998; Hauk et al., 2011; Phillips, Rugg, & Friston, 2002; Zetter, Iivanainen, Stenroos, & Parkkonen, 2018). Here, we will extend these findings by presenting three metrics for both PSFs and CTFs, as well as five source estimation methods and two sensor configurations. We will first describe the framework for linear resolution analysis of EEG/MEG source estimation in more detail. The central concept is the resolution matrix, which contains both PSFs and CTFs. The most popular source estimation methods will be described in this framework, namely different types of L2-minimum-norm estimation and spatial filters such as beamformers. We will then compare five of these methods with respect to spatial resolution metrics capturing localization accuracy, spatial extent and relative amplitude.

## Theory and Methods

### Linearity and the superposition principle

Linear transformations, by definition, have the property that the output of any sum of inputs is just the sum of the outputs for the individual inputs (“ superposition principle”). More formally, a transformation **T** is linear if for any arbitrary input signals A and B and scalar values *a* and *b* the following is true:

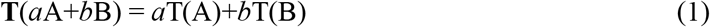

In the case of EEG/MEG source estimation, A and B can be dipolar current sources with unit strength, *a* and *b* source strengths, and the transformation **T** the inverse matrix that turns the measured EEG/MEG data into current distributions in the brain.

This means that we can characterize the properties of the transformation on the basis of the transformations of individual point sources with unit strength, or their point-spread functions (PSFs). This is a standard procedure for linear systems, whether in the context of Hifi systems, microscopes, or source estimation (https://en.wikipedia.org/wiki/Point_spread_function, illustrated in Figure 1). Figure 1A shows example point-spread functions for a microscope, i.e. the degree of blurring for two circular objects due to the microscope’s point-spread function. The same principle applies to EEG/MEG topographies in signal space (Figure 1B). Here, it is also shown how localization accuracy, spatial extent and relative amplitude are all relevant to describe spatial resolution. Figure 1C demonstrates how the superposition principle applies to source estimation (here L2-minimum-norm estimates).

**Figure 1:**
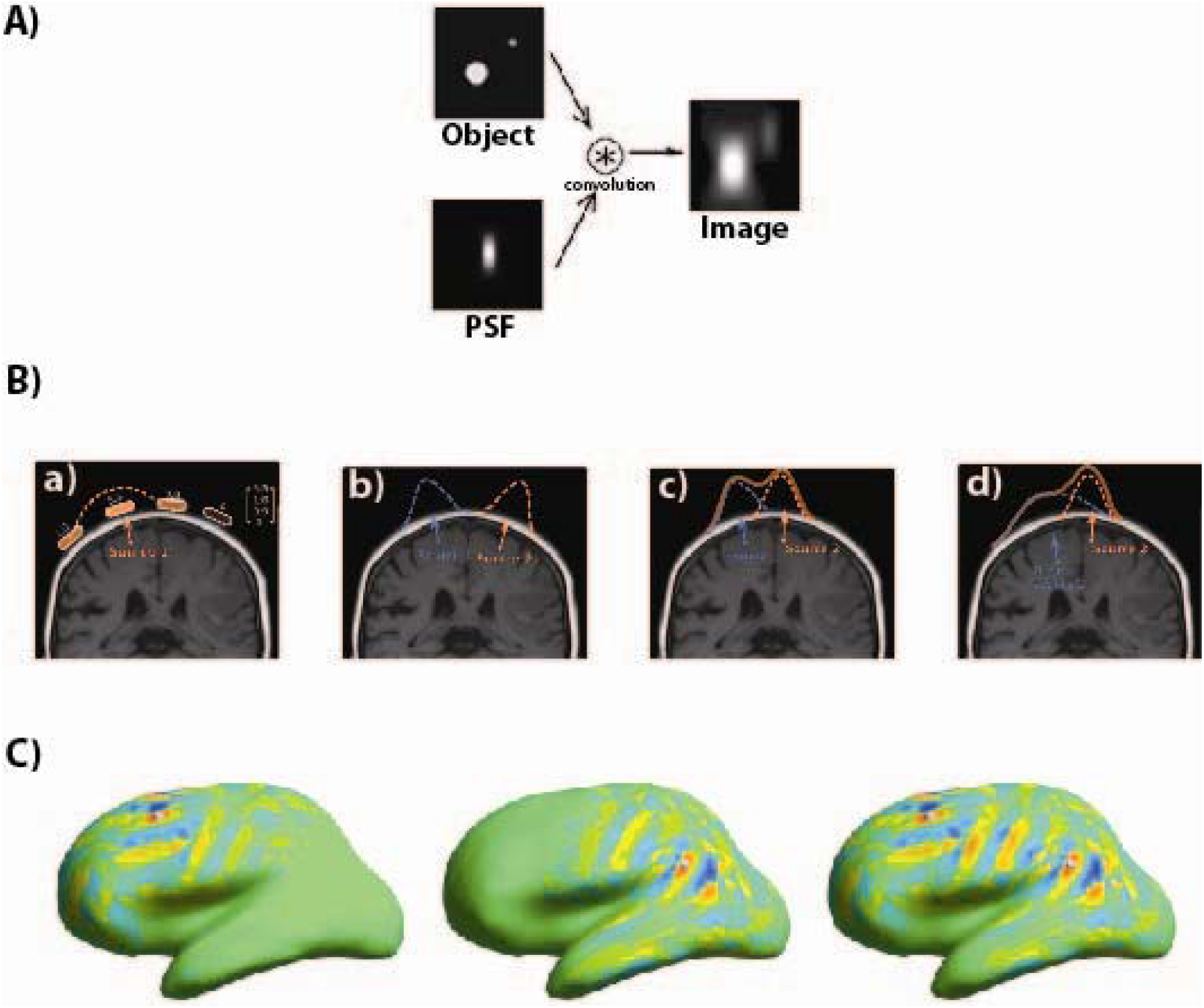
Illustration of the superposition principle for linear data transformation. **A)** A hypothetical microscope “blurs” point-like objects according to the point-spread function (PSF, bottom left). For two real objects (top left), the output image (right) is the real image convolved (or “blurred”) with the PSF. **B)** Illustration of the superposition principle for the EEG forward problem. a) A point source (current “dipole”) produces a characteristic voltage distribution, or topography, on the scalp. This topography is represented by a column vector (top right). b) The topographies of two separate sources add up at the scalp. If they are sufficiently distant from each other, their topographies hardly overlap. c) As the sources move closer together, their topographies begin to overlap. d) For close-by sources with different depths, the topography of the superficial source may overshadow the topography of the deep one. The Gaussian-shaped EEG topographies for radial dipoles in this example were chosen for simplicity, and are only a very coarse approximation of reality. a and b are from https://en.wikipedia.org/wiki/Point-spread_function). **C)** The superposition principle for linear distributed source estimates. The source distributions for two point sources (PSFs) are shown in panels 1 and 2, respectively (L2-MNE, MEG only). The locations of the point sources are indicated by two small balls on the inflated cortical surface. The source distribution for both sources together is just the sum of their individual PSFs (panel 3). Red and blue colours indicates out- and in-going source currents with respect to the cortical surface, respectively.

### Resolution Matrix

The resolution matrix is a useful tool for the characterization of spatial resolution for linear estimators (Dale & Sereno, 1993; Menke, 1989). It describes the relationship between the true unknown sources **s** and the estimated sources **ŝ**. Thus, while we won’t be able to know the true sources **s**, the resolution matrix can help us estimate how far we are away from the truth, and for which source locations we may at least be close.

The logic behind the resolution matrix is as follows: We know that our measured data are the result of the forward solution for the source distribution, and our linear source estimate is the result of the linear estimator applied to our measured data. If we combine the forward solution and the inverse estimator together into one transformation, we directly transform the unknown true source distribution into its estimate. Ideally, this transformation should be the identity, i.e. our estimate should be exactly the true source distribution. We know that we cannot achieve this due to fundamental physical and mathematical limitations of the inverse problem, but an inspection of this transformation can tell us how we close we are. This transformation is given by the resolution matrix, which will now be derived more formally.

The relationship between the source distribution **s** (vector mx1) and the data in n channels **d** (nx1) can be written in matrix form

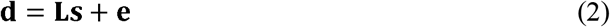

where **L** is the so-called leadfield matrix which contains information about head geometry (shape and conductivities), measurement configuration (sensor types and position), and the physics of signal generation (quasi-static approximation of Maxwell’s equation (Sarvas, 1987)), and **e** is the error term. This forward problem is necessarily linear (illustrated in Figure 1B).

Source estimation methods do not have to be linear, but we can try to find an optimal linear matrix **G** (optimal in a way to be defined), such that we obtain an estimate **s** for the source distribution

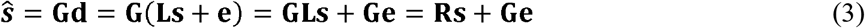

where we have made use of the previous equation. The resolution matrix **R** provides the desired relationship between true and estimated sources, and the last term takes into account how noise is transformed from signal space into source space. This will be relevant for regularization procedures later.

### Point-spread and cross-talk functions (PSFs and CTFs)

As illustrated in Figure 1, linear transformations can be meaningfully evaluated for point sources. PSFs and CTFs address two aspects of linear source estimation methods: 1) How does the method spread the activity of a point source with unit strength to other locations (PSF), and 2) how is the result of the method for one particular source affected by other sources (CTF)? In order to derive PSFs and CTFs more formally, it is useful to break down equation (3) for individual elements **s** of and **ŝ**.

In order to obtain the PSF for source i, **PSF**_*i*_, we need to obtain the source estimate for the topography in sensor space produced by source i with unit strength. This topography is the i-th column of the leadfield matrix, i.e. **L**_.i_. Therefore:

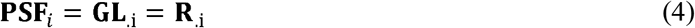

In other words, the PSF for source i is the i-th column of the resolution matrix **R**. This also shows that **PSF**_*i*_ is a weighted sum of the columns of the inverse matrix **G**, with the weights given by **L**_.i_.

The CTF for source i, **CTF**_*i*_, is obtained by applying the linear estimator for source i to the topographies of all possible point sources with unit strength. The estimator for source i is the i-th row of the inverse matrix **G** i.e. **G**_*i*._. The topographies are all arranged as columns of the leadfield matrix **L**. Thus

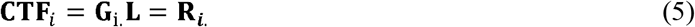

In other words, the CTF for source i is the i-th row of the resolution matrix. It also shows that **CTF**_*i*_ is the weighted sum of the rows of the leadfield matrix, with the weights given by **G**_i._. Let us put the formulae for PSFs and CTFs next to each other:

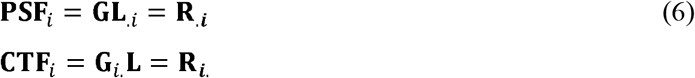

As stated above, PSFs are the columns of the resolution matrix and linear combination of columns of the inverse matrix **G**, and CTFs are the rows of the resolution matrix **R** and linear combinations of the rows of the leadfield matrix **L**. In general, the PSF and CTF for a source i will have different shapes, which is important for the comparison of different source estimation methods. However, the resolution matrix for the unweighted L2-minimum-norm estimate is symmetric, and for this method PSFs and CTFs can be represented by the same distributions.

While the leadfield matrix is determined by our forward model, the inverse matrix can be chosen arbitrarily in order to optimise our PSFs and CTFs. But whatever we do, linear algebra poses some fundamental limits (Menke, 1989):

1. A distribution that cannot be approximated by any combination of the rows of the leadfield matrix cannot be a CTF.
2. The number of columns in our estimator is the number of recording channels N_C_. We can therefore get at most N_C_ ideal PSFs, but this does not guarantee that the remaining N_S_-N_C_ PSFs are ideal (or even close) as well.
3. Because **G** appears in both formulae, we cannot optimise PSFs and CTFs independently of each other.

This implies that every optimisation of PSFs and CTFs will require a trade-off between different resolution criteria. We will address this issue using quantitative resolution metrics below.

### Noise and regularisation

Source estimation attempts to determine the brain sources that generated the measured signals. In this process, it tries to suppress signals that are likely to originate from outside the brain, i.e. it should minimise the term **Ge** in equation (3). This process if often called regularisation of the source estimates. In short, we accept some loss of spatial resolution and/or goodness-of-fit in return for a solution that is less effected by noise, which usually results in smoother source distributions (Backus & Gilbert, 1968; Bertero, De Mol, & Pike, 1988; Menke, 1989). The degree of regularisation is controlled by the regularisation parameter, often denoted as λ (lambda). The larger this parameter, the lower the spatial resolution and/or goodness-of-fit, and the higher the smoothness of the source distribution.

The details of regularisation procedures are not within the scope of this paper. However, for a fair methods comparison is important to choose the regularisation parameter in similar ways for all methods. This can for example be accomplished via the SNR of the data. Furthermore, the regularisation parameter determines the effect of the noise covariance matrix on the inverse matrix. If λ = 0, the noise covariance matrix does not matter at all. The larger λ, the more it matters. Thus, in order to compare methods, and in order to generalise results to other data sets, the structure and regularisation of the noise covariance matrix has to be similar.

### Linear source estimation methods

Inverse matrices and spatial filters can be obtained in a number of different ways. Here, we will only briefly mention the most popular ones. A more detailed description, analysis and comparison of these methods is not within the scope of this paper. Our general approach, i.e. the use of the resolution matrix and resolution metrics to evaluate spatial resolution, is applicable to any other linear method as well.

### L2-Minimum-Norm-Type Methods

A common expression for the unweighted L2-MNE is

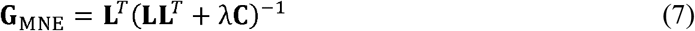

where *λ* is the regularization parameter and **C** the noise covariance matrix. The resolution matrix in this case is

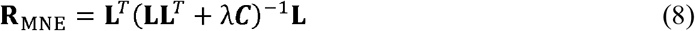

which is a symmetric matrix, and therefore PSFs and CTFs for elements *i* are the same. L2-MNE produces the resolution matrix which is closest to the ideal case, i.e. the identity matrix, in the least-squares sense (Menke, 1989). This property suggests that L2-MNE provides an optimal trade-off of different spatial resolution criteria. However, it has often been pointed out that L2-MNE shows a strong tendency to localize sources too close to the sensors. Two strategies have been proposed to alleviate this: Depth-weighting (Lin et al., 2006) and (noise-)normalization (Dale et al., 2000; Pascual-Marqui, 2002).

Depth-weighting can be implemented by introducing a diagonal weighting matrix **D** into equation 7:

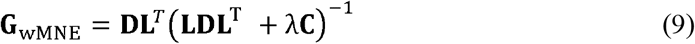

The values in **D** can be chosen to “boost” the impact of deep sources, e.g. based on the vector norms of the leadfield columns (topographies of deeper sources tend to have smaller norms). Note that the same weighting strategy can be chosen to include a priori information about source locations, e.g. from fMRI data: the more we boost activation of certain sources via **D**, the more we bias our source distribution towards these sources, possibly even when these sources are actually not active (Ahlfors & Simpson, 2004). Thus, depth-weighting can also be interpreted as using a priori information to bias the source estimate towards deeper sources.

Another way to change the localization properties of the L2-MNE estimator is (noise-)normalization (Dale et al., 2000; Pascual-Marqui, 2002). In this case, the L2-MNE matrix is multiplied with a diagonal weighting matrix **W**:

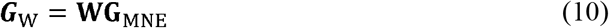

The resolution matrix then changes accordingly:

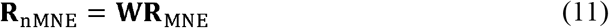

Thus, we may find a **W** that improves some desirable properties of the resolution matrix. Because the **W** matrices are diagonal, each row *i* of the MNE resolution matrix is scaled by the factor *W_ii_*. As a consequence, the shapes of the CTFs (rows of **R**) do not change (Hauk et al., 2011). Only the shape of PSFs (columns of **R**), and therefore potentially locations of peaks and their spatial extensions, is affected by this normalization procedure.

The two most popular methods of this type are dynamic statistical parametric mapping (dSPM) and standardized low-resolution electromagnetic tomography (sLORETA):

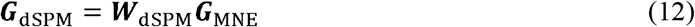

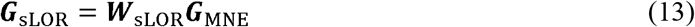

For dSPM, the normalization matrix contains the minimum norm estimates of the noise at each source (Dale et al., 2000), derived from the noise covariance matrix, i.e.

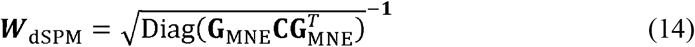

For sLORETA (Pascual-Marqui, 2002), the normalization uses the diagonal of the MNE resolution matrix

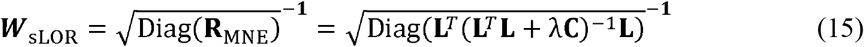

It has been shown that dSPM and sLORETA have better peak localization performance for PSFs than L2-MNE (Hauk et al., 2011), and even that sLORETA has zero dipole localization error (Pascual-Marqui, 2002) under ideal circumstances. However, localization performance does not allow similar conclusions about other aspects of PSFs (e.g. spatial extent, local extrema etc.), or about the shape of CTFs (Hauk et al., 2011). We do not evaluate eLORETA, also a method with zero peak localization error, in the present study (Pascual-Marqui et al., 2018). This should be done in a future study.

### Beamforming

We stated above that linear inverse methods can be interpreted as “spatial filters” or “software lenses” that attempt to estimate activity from sources of interest, while suppressing interference from all other possible sources. These spatial filters can be arranged as rows of an inverse matrix to provide the whole source distribution. In this logic, the ideal inverse matrix should yield a resolution matrix as close as possible to the identity matrix. Thus, an individual spatial filter’s CTF should be as close as possible to the corresponding row of the identity matrix. If closeness to the identify matrix is required in the least-squares sense, this yields the L2-MNE solution (Backus & Gilbert, 1968; Menke, 1989).

Beamformers approach this problem differently. They start from the requirement that a spatial filter for source i should be sensitive to activity from this source, i.e. it should yield the value 1 if this source is active with unit strength (unit gain beamformer) (Van Veen et al., 1997):

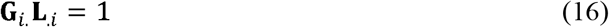

At the same time, the beamformer attempts to suppress other activity. In contrast to minimum-norm type methods, it does not do this by modelling other sources via the leadfield matrix, but by assuming that they are captured by the data covariance matrix **C**_D_. Note that this matrix contains the noise covariance as well as possible signal covariance from other sources of interest. The requirement to suppress the effect of other activity on the estimate for source i can thus be written as

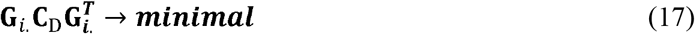

This results in the unit-gain linearly constrained minimum variance beamformer:

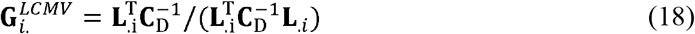

Variations exist that optimise beamformers for pairs of dipolar sources (Brookes et al., 2007) and for time- and frequency-dependent beamforming (Dalal et al., 2008; Woolrich et al., 2013).

Because the estimator is a vector that is applied to the data, we can use the concept of CTFs to describe its spatial resolution. In principle, the superposition principle holds, since the contributions of different sources add up. However, the situation is more complicated in this case: Because we are using the data covariance matrix, the estimator depends on the sources of interest. Whenever the data change, so does the estimator. This is why beamformers are often called “adaptive spatial filters” (Sekihara & Nagarajan, 2008) – they adapt to the data. In contrast, minimum-norm-type estimators are “static”, since they are based only on the leadfield and noise covariance matrices (assuming that the noise structure does not change during the experiment). This affects the generalisability of any resolution analysis: While the result for linear minimum-norm type methods apply to other data sets with similar measurement configurations, head models and noise structures, the results for beamformers are only valid under those conditions that are represented in the data covariance matrix. Thus, while beamformers yield a linear transformation of the data, their adaptivity to data makes their PSFs and CTFs hard to generalise to different data sets.

Therefore, we opted for a non-adaptive spatial filter in this study, using the noise covariance matrix instead of the data covariance matrix in equation 28:

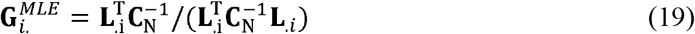

The data covariance matrix in any given experiment is the noise covariance matrix plus the signal covariance (assuming additivity of noise and signal, which is a common assumption). If we were going to estimate the sources in our baseline interval, then this filter could be interpreted as a beamformer, since in this case **C**_N_ would represent the activity due to the sources of interest. However, equation 19 also describes the maximum-likelihood estimator for the strength of a single dipole at position *i* (Lütkenhöner, 1998a). Therefore, we will refer to this estimator as a dipole filter (DF).

Alternatively, we could have simulated a data covariance matrix under the assumption that all sources in the model are randomly activated, with uncorrelated time courses in the source vector **s**_N_(*t*), following a uniform Gaussian distribution (independent identically-distributed sources, i.i.d). In that case, the data covariance matrix would be

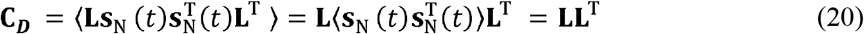

If we insert this into equation 19, we obtain the L2-MNE estimator with a different scaling; this is a natural outcome, as the assumed i.i.d source activity matches the implicit prior of L2-MNE. While the scaling will affect the PSFs, it will not affect the shape of the CTFs, and the CTFs results for L2-MNE will also apply to this type of beamformer. Thus, we only used the spatial filter of equation 19. A more detailed evaluation of beamformers will be left for future studies.

### Resolution Metrics

We stated above that a meaningful resolution analysis of EEG/MEG source estimation methods should include metrics for the three resolution categories: localization accuracy, spatial extend and relative amplitude. Here we provide a table (Table 1) with possible metrics, most of which have already been used in previous studies (but not necessarily together). We will use a subset of those in our Results section.

**Table 1:**
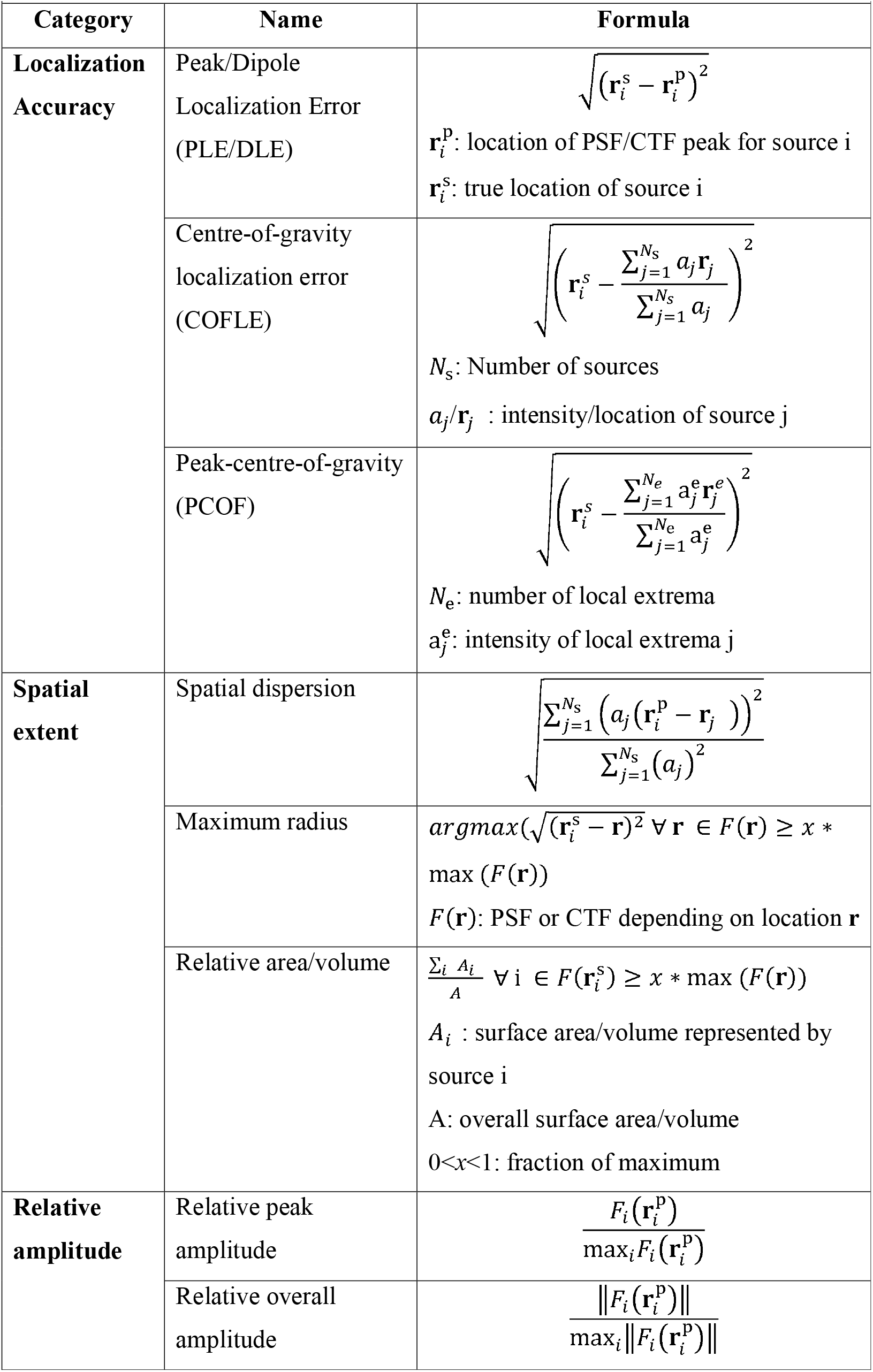
Spatial resolution metrics.

### Simulation set-up

We computed resolution matrices and resolution metrics for the open EEG/MEG data provided by Wakeman and Henson (2015). It consists of 16 subjects from which simultaneous EEG and MEG were recorded in an Elekta Neuromag Vectorview scanner (70 electrodes, 102 magnetometers, 204 planar gradiometers), in addition to MRI, fMRI and DTI data. Details of the data acquisition parameters are available in the previous publication (Wakeman & Henson, 2015). In the following, we will only summarise the most important parameters for our purposes.

For our simulations, we only required the sensor configurations, structural MRI images, as well as noise covariance matrices (details below). Pre-processing of EEG/MEG and MRI data, including source estimation, was carried out in MNE-Python software. Resolution analysis was performed in Python, making use of MNE-Python functionality where appropriate. The software used for our resolution analysis is available online (https://github.com/olafhauk/EEGMEGResolutionAtlas). The noise covariance matrices for each data set were computed concatenating pre-stimulus baseline intervals of 200 ms duration before stimulus onset. Stimuli in this experiment were pictures of faces and scrambled faces. The noise covariance matrix was estimated in empirical mode in MNE-python, and regularized using a rank-dependent Tikhonov approach with regularization parameter 0.2 for all sensor types separately. For regularization of the inverse estimators, the default signal-to-noise ratio in the MNE software was used (SNR=3). This value is representative for many event-related EEG/MEG studies.

High-resolution structural T1-weighted MRI images were acquired in a 3T Siemens Tim Trio (Siemens, Erlangen, Germany) scanner at the CBU using a MPRAGE sequence, and processed using MNE-Python software, which makes use of PySurfer. This included the segmentation of the cortical surface as source space, and the creation of a three-compartment boundary element model (BEM) for the forward solution. MEG sensor configurations and MRI images were coregistered based on the matching of about 50-100 digitized locations on the scalp surface with the reconstructed scalp surface from the structural MRI.

The source space consisted of the cortical surface with 4098 vertices per hemisphere (octahedron spacing 6). BEMs were created using linear collocation with 5120 triangles per surface. The surfaces represented the inner skull, outer skull and outer skin, respectively.

The forward solution was originally computed with free orientation constraint, i.e. three source orientations per location. At the stage of computing the inverse operators, sources were constrained to be approximately perpendicular to the cortical surface, with the variance of the tangential component being 0.2 of the variance of the perpendicular component (“loose orientation constraint”).

The computation of all inverse matrices was based on standard functionality in MNE-Python. Inverse matrices and leadfield matrices were extracted from the inverse operators and forward solutions, respectively, in order to compute the resolution matrix and the resolution metrics for each individual subject. Those results were then morphed to a standard brain (Freesurfer’s “fsaverage”), and averaged across subjects. Results will be shown for the average brain. We do not expect large qualitative differences between left and right brain hemispheres, and therefore only the left hemispheres will be shown.

### Results

In a first step, we compared an EEG+MEG with an MEG-one configuration similar to Molins et al.(2008). Figure 2 shows the resolution metrics peak localization error (PLE), spatial deviation (SD), and relative overall amplitude (ROA) for unweighted L2-MNE. Results are shown for combined EEG plus MEG, for MEG only, and for the difference of the two measurement configurations. Note that for the MNE estimator the resolution matrix is symmetric, and in this case the results apply to both PSFs and CTFs. Figure 2 A and B show that for locations that are further away from the sensors, e.g. in the Sylvian fissure, all resolution metrics indicate lower resolution: PLE and SD increase and ROA decreases with source depth. Performance is generally better for combined EEG+MEG (A) than for MEG alone (B), which is confirmed by the subtractions (C): Better performance for EEG+MEG results in blue colors for PLE and SD and in red colors for ROA.

**Figure 2:**
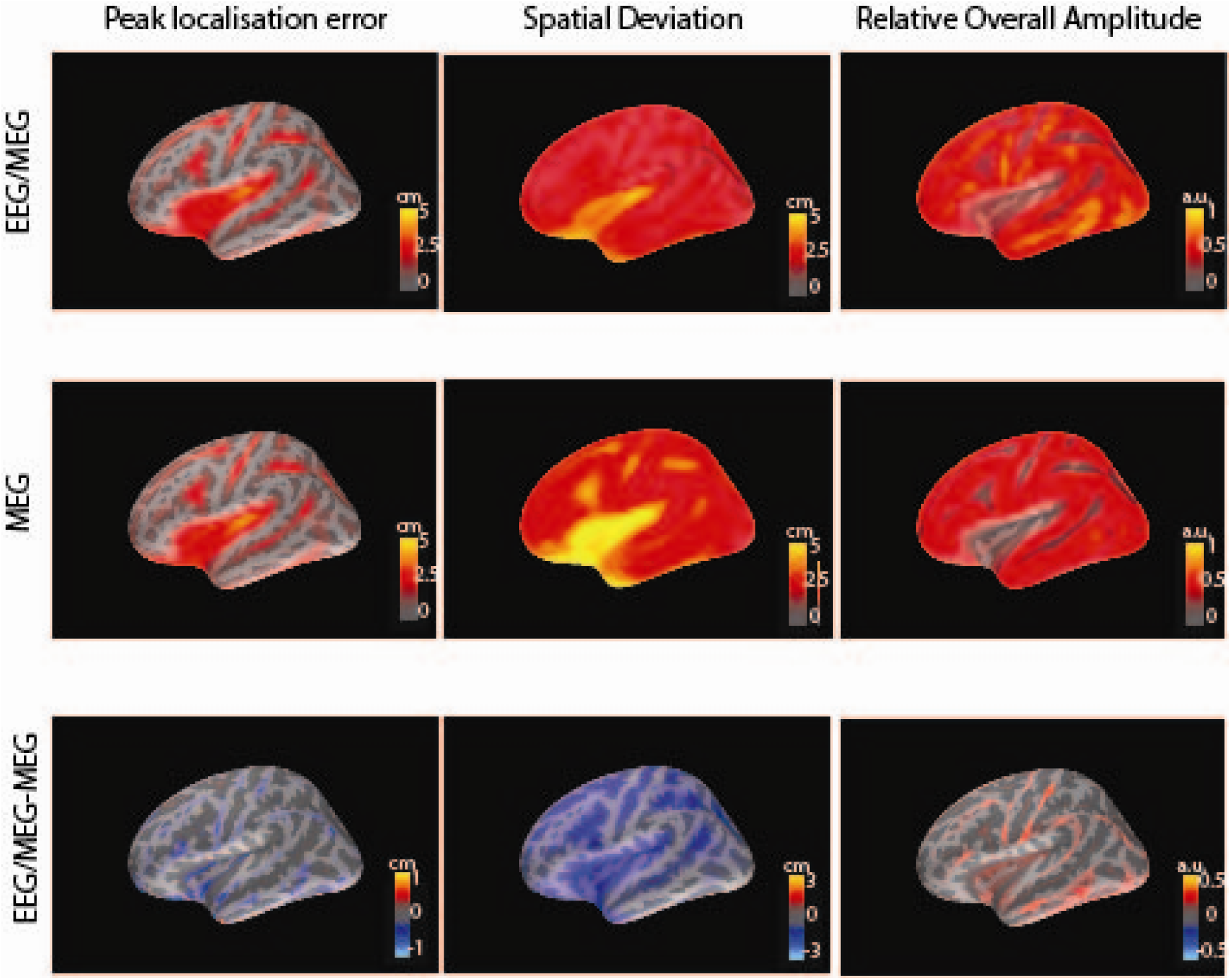
Comparison of EEG/MEG and MEG sensor configurations for L2-MNE estimation. Peak localisation error, spatial deviation and relative overall amplitude for point-spread and cross-talk functions (PSFs/CTFs) are shown as distributions across the left cortical hemisphere for combined EEG+MEG (A), MEG (B), and the subtraction of MEG from EEG+MEG (C). Note that for L2-

In the following figures 3–5 we present methods comparisons for the combination of EEG and MEG, i.e. for the best-case scenario. We extend previous results by Hauk et al. (2011), who used MEG-only for a smaller number of methods. Figure 3 shows the PLE distributions for PSFs and CTFs for several methods: unweighted L2-MNE, depth-weighted (dw) L2-MNE, dSPM, sLORETA, and the DF. Rows 1 and 3 show distributions for the individual methods, and rows 2 and 4 the subtractions with respect to L2-MNE. For PSFs, sLORETA and MLE-BF exhibit the predicted zero PLE. All methods show some improvement over L2-MNE with respect to PLE, especially for deeper source locations. For CTFs, the performance of all methods is very similar. This was predicted for L2-MNE, dSPM and sLORETA, as their CTFs have the same shape. This result demonstrates that the localization properties of PSFs are easier to manipulate than the crosstalk in CTFs.

**Figure 3:**
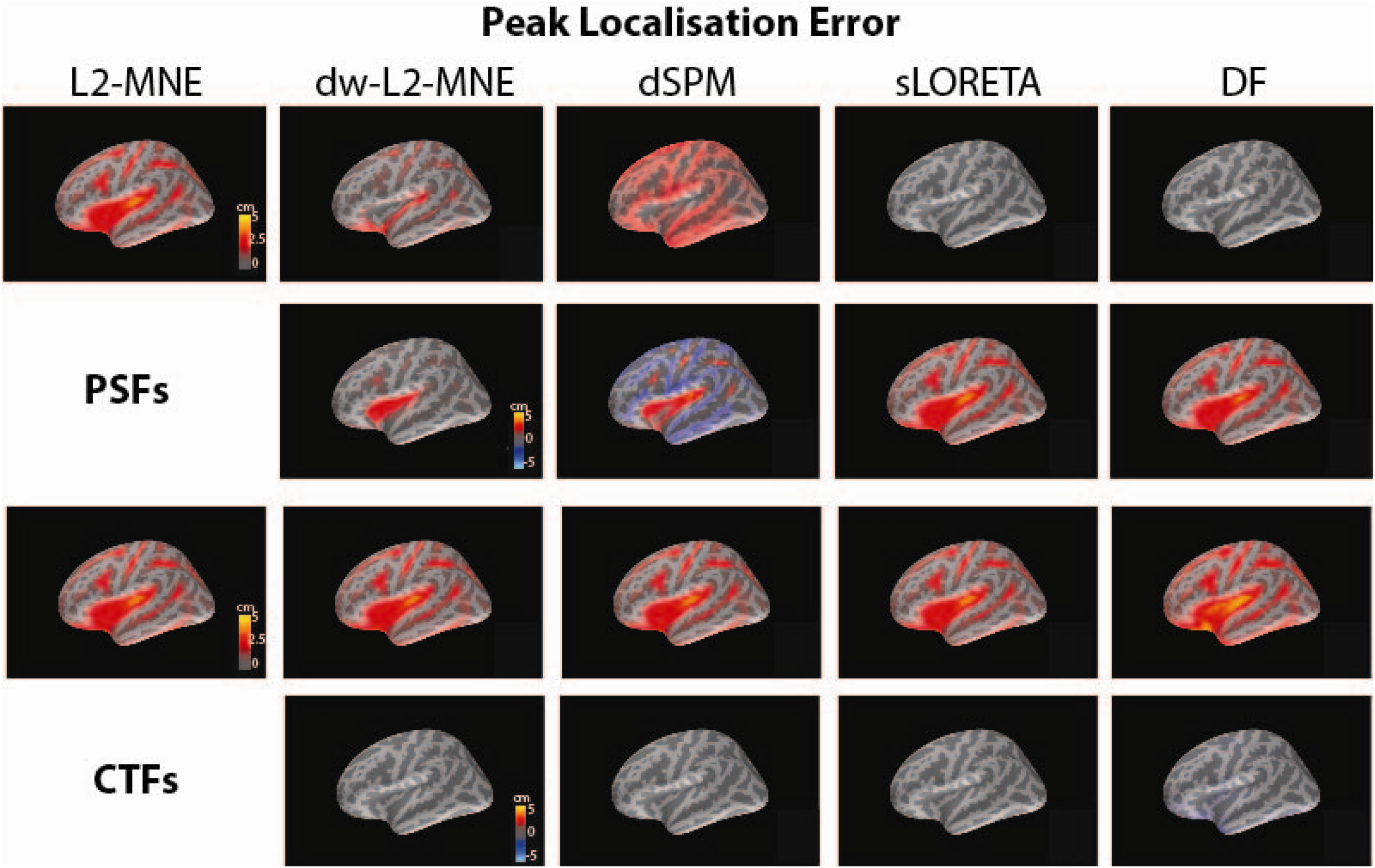
Comparison of PSFs and CTFs with respect to peak localisation error. Peak localisation error is shown for PSFs (top) and CTFs (bottom) and five linear source estimation methods (columns) as distributions across the left cortical hemisphere. Rows 1 and 3 show distributions for individual methods, and rows 2 and 4 their subtractions from L2-MNE. Each row has the same colour scaling. PSF/CTF: point-spread/cross-talk function; MNE: minimum-norm estimator; dw: depth-weighted; dSPM: dynamic statistical parametric mapping; sLORETA: standardised low resolution electromagnetic tomography; DF: dipole filter.

**Figure 4:**
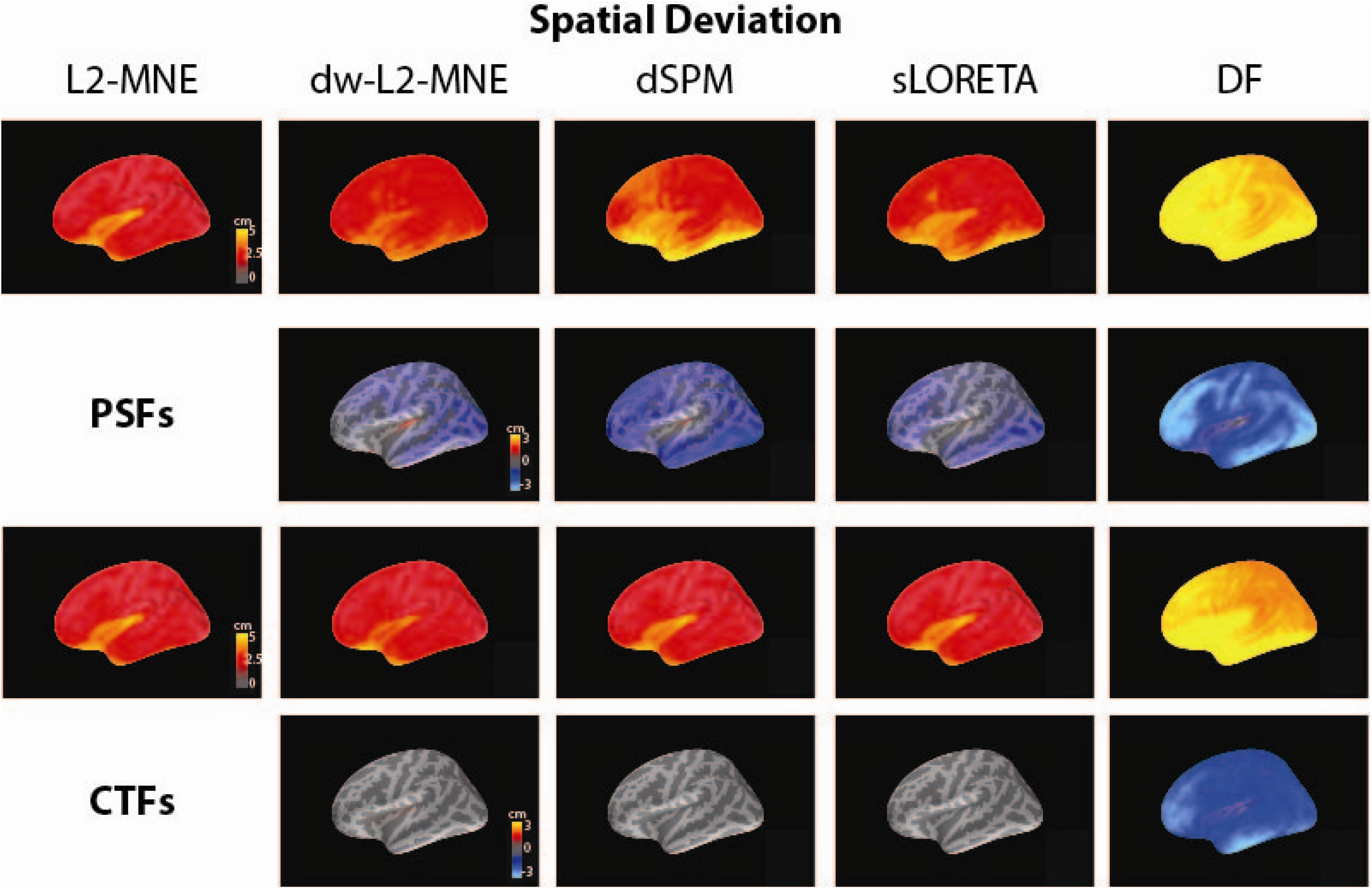
Comparison of PSFs and CTFs with respect to spatial deviation. Spatial deviation is shown for PSFs (top) and CTFs (bottom) and five linear source estimation methods (columns) as distributions across the left cortical hemisphere. Rows 1 and 3 show distributions for individual methods, and rows 2 and 4 their subtractions from L2-MNE. Each row has the same colour scaling. Abbreviations as for Figure 3.

**Figure 5:**
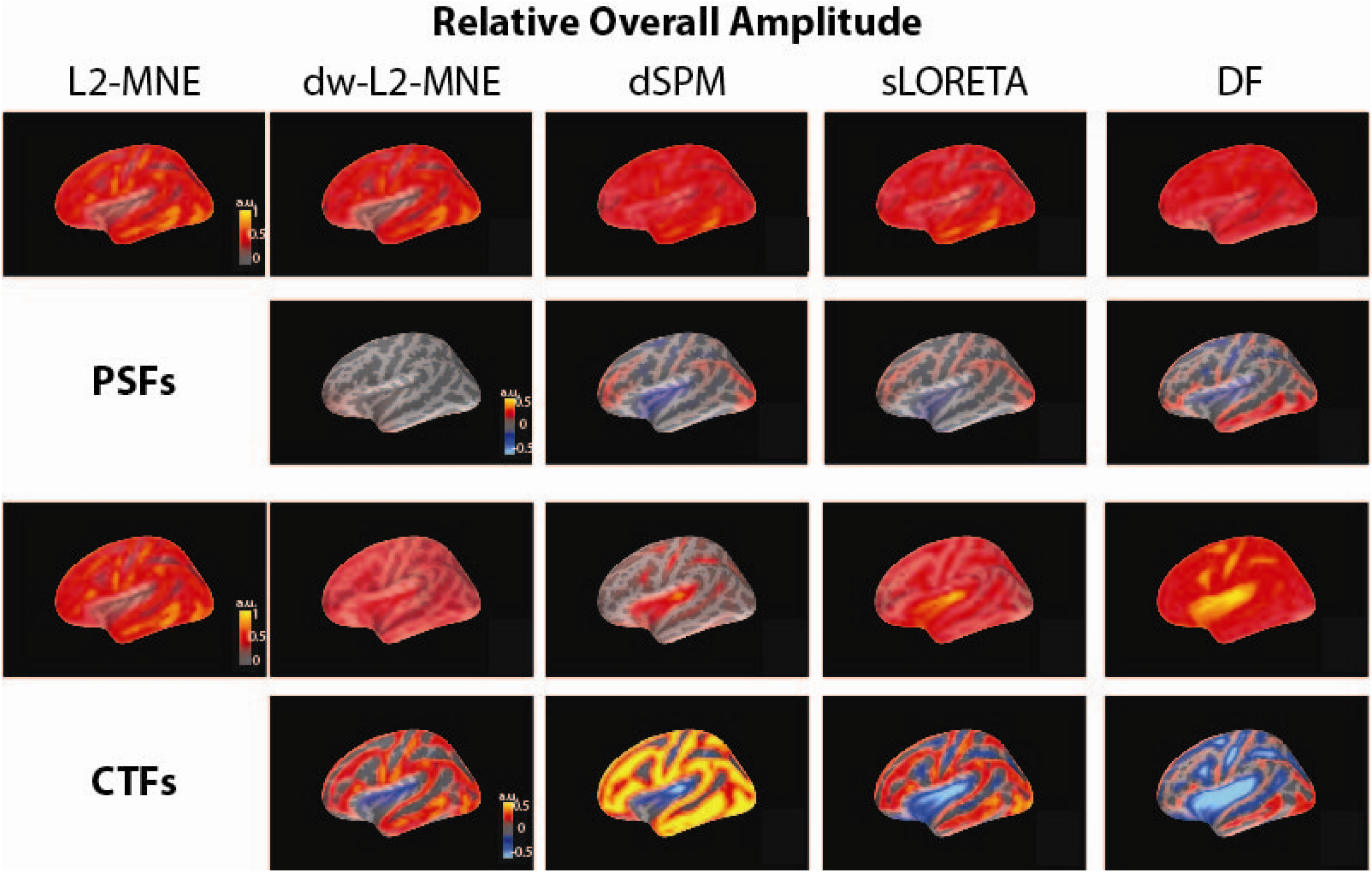
Comparison of PSFs and CTFs with respect to relative overall amplitude. Relative overall amplitude is shown for PSFs (top) and CTFs (bottom) and five linear source estimation methods (columns) as distributions across the left cortical hemisphere. Rows 1 and 3 show distributions for individual methods, and rows 2 and 4 their subtractions from L2-MNE. Each row has the same colour scaling. Abbreviations as for Figure 3.

Figure 4 shows the analogous results for spatial deviation. In contrast to PLE, SD increases with source depth for all methods, indicating a loss of spatial resolution. This is the case for both PSFs and CTFs. The subtractions for PSFs show that all methods perform worse than L2-MNE with respect to SD. As for PLE, the SD distributions for CTFs are the same for L2-MNE, dSPM and sLORETA, and they are very similar to dw-L2-MNE. However, DF clearly stands out as the worst performer with respect to SD.

Figure 5 presents the results for relative overall amplitude. For PSFs the distributions are similar, with decreasing amplitudes for deeper sources. The subtractions do not reveal a clear winner: While L2-MNE performs better in superficial locations, dSPM, sLORETA and MLE-BF perform better for deeper sources. CTFs show a similar yet more complex pattern of results. L2-MNE shows largest values in superficial regions, while for the other methods the largest values appear at deeper locations. As for PSFs, L2-MNE’s advantage for superficial locations and worse performance for deeper sources is confirmed in the subtractions, although in this case DF shows generally better performance.

## Discussion

We provided a framework and the basic methodology to evaluate spatial resolution for linear EEG/MEG source estimation methods. Spatial resolution cannot be quantified by a single number, and the choice of appropriate resolution metrics ultimately depends on the goal of the experimenter. Here, we showed that in order to draw inferences for cases involving multiple sources, at least three classes of resolution metrics should be taken into account to evaluate point-spread and cross-talk functions (PSFs/CTFs): localization accuracy, spatial extent, and relative amplitude. In our simulation we used on example for each of these classes, namely peak localization error, spatial deviation, and relative overall amplitude. Other resolution metrics can easily be implemented in the future.

Spatial resolution depends on a large range of variables (sensor configurations, source locations, SNR, head model, etc.), which may vary significantly across studies, or even across different analyses within one study. It is therefore essential to provide researchers with tools to evaluate the spatial resolution of their analysis setup at hand. So far, existing EEG/MEG software packages only provide very limited functionality for the evaluation of spatial resolution. We here present results for a common measurement configuration and parameter settings, in order to illustrate the usefulness of the approach and address some important issues with respect to the combination of different sensor types and methods comparison. Other parameters, e.g. novel on-scalp MEG sensor configurations or head models, should be investigated in the future. (Boto et al., 2018; Iivanainen et al., 2017; Tierney et al., 2019).

We argued that spatial resolution should not just be evaluated on PSFs, but also for CTFs. PSFs are more flexible in so far as they can be arbitrarily designed for a certain number of sources (smaller or equal to the number of sensors). This feature can be exploited for example to include a priori information, such as to increase localization accuracy in regions where sources are expected, at the expense of lower localization accuracy in other regions (Ahlfors & Simpson, 2004). CTFs, on the other hand, are strongly constrained by the leadfields. Thus, while we are able to design an estimator that will produce a peak in the PSF for this source at the correct location, in reality the source estimate at this location may still receive a large amount of leakage from other brain regions. We therefore recommend to evaluate the spatial resolution of linear methods using PSFs as well as CTFs.

This study focused on linear non-adaptive methods. For those methods, point-spread and cross-talk functions (PSFs and CTFs) yield generalizable results due to the superposition principle. Thus, no matter which constraints were used to design the corresponding inverse matrices, PSFs and CTFs provide an assumption-free evaluation of their spatial resolution that can be compared across methods. Sparsity (i.e. the presence of a small number of focal sources) is sometimes used as a nonlinear constraint, e.g. in L1-minimum-norm-type methods (Huang et al., 2014; Owen, Wipf, Attias, Sekihara, & Nagarajan, 2012; Uutela, Hamalainen, & Somersalo, 1999), or multiple sparse priors (Friston et al., 2008). These nonlinear methods are hard to evaluate in a generalizable way, and their results strongly depend on the validity of the underlying assumptions (Krishnaswamy et al., 2017). For example, perfect performance for point sources in isolation does not directly imply perfect performance for multiple simultaneously active sources (Grave de Peralta, Hauk, & Gonzalez, 2009). The simulation setups for non-linear methods have to be chosen and evaluated carefully in order to represent source scenarios that can be realistically assume to underlie measure brain activity in certain types of experiments. Similar setups should work for beamformers whose filters depend on the data covariance matrix, i.e. on sources of interest.

Recent EEG/MEG research has increasingly focused on patterns of brain activations (Kietzmann et al., 2019; Stokes, Wolff, & Spaak, 2015). While any pattern can be modelled as the sum of point sources, it is not obvious to predict how an activity pattern in one ROI will affect patterns in other ROIs. This will depend on features of these patterns, e.g. their spatial frequency, homogeneity, etc. This requires a statistical approach, i.e. the simulation of a large number of representative patterns in one ROI and their effects in other ROIs. This should be investigated in future studies.

The results of the present study confirm and generalize those obtained in previous studies (Hauk et al., 2011; Molins et al., 2008). We could show that combining EEG and MEG improves spatial resolution, especially with respect to spatial extent (Molins et al., 2008). While in our study we cannot rule out that this is merely to an increase in the number of sensors, a previous study has shown that EEG indeed adds independent information to MEG recordings (Molins et al., 2008). This reflects the fact that the leadfields for EEG and MEG sensors are more similar within than across their measurement modality, i.e. they are sensitive to different aspects of the current distribution (Goldenholz et al., 2009; Henson, Mouchlianitis, & Friston, 2009; Malmivuo, 2012). We here extended this finding to the resolution metric relative amplitude.

Our method comparison confirms previous results that EEG//MEG spatial resolution is inherently limited, and different methods can only make different compromises among different resolution criteria (Hauk et al., 2011). For example, (noise-)normalization improves the localization accuracy of L2-MNE, but at the cost of increased spatial extent and worse relative amplitude. Our results extend those findings to depth-weighed minimum-norm estimation and a non-adaptive spatial filter, as well as to combined EEG and MEG measurement configurations.

It is interesting to note that our simple dipole filter had zero peak localization error, a property it shares with sLORETA. However, it shows worse performance with respect to spatial extent than the other methods. This is due to the fact that this spatial filter for a specific source location does not attempt to suppress the cross-talk from any other source in the model (i.e. the leadfield), but instead attempts to suppress the signals captured in the noise covariance matrix. For beamformers, the data covariance matrix would contain contributions from the signals of interest plus the noise covariance. In cases where the signal of interest is generated by a few focal sources, the rejection of their signals may dominate the shapes of the corresponding CTFs. It has been argued previously that “this is the reason why adaptive spatial filters have a strange-looking beam response” (another term for cross-talk function; Sekihara and Nagarajan, p. 42).

While we observed striking differences between methods with respect to their PSFs, these differences were much smaller for CTFs. This reinforces our argument from above that linear methods should be evaluated both on the basis of PSFs and CTFs. On balance, we conclude that if no strong prior information exists then L2-MNE is the best compromise of different resolution criteria. In any case, our approach to define multiple resolution metrics based on PSFs and CTFs will allow researchers to make their own compromises. In the future, we plan to implement resolution analysis tools to common EEG/MEG analysis software packages such as MNE-Python.

## Acknowledgments

This work was supported by the Medical Research Council UK (SUAG019 RG91365).

## References

Ahlfors, S. P., & Simpson, G. V. (2004). Geometrical interpretation of fMRI-guided MEG/EEG inverse estimates. Neuroimage, 22(1), 323–332. doi:10.1016/j.neuroimage.2003.12.044 S1053811904000199 [pii]

Backus, G. E., & Gilbert, J. F. (1968). The resolving power of gross earth data. Geophysical Journal of the Royal Astronomical Society, 16, 169–205.

Baillet, S., Friston, K., & Oostenveld, R. (2011). Academic software applications for electromagnetic brain mapping using MEG and EEG. Comput Intell Neurosci, 2011, 972050. doi:10.1155/2011/972050

Baillet, S., Mosher, J. C., & Leahy, R. M. (2001). Electromagnetic brain mapping. IEEE Signal Processing Magazine, 18(6), 14–30.

Bertero, M., De Mol, C., & Pike, E. R. (1988). Linear inverse problems with discrete data: II. Stability and regularisation. Inverse Problems, 4(3), 573–594.

Boto, E., Holmes, N., Leggett, J., Roberts, G., Shah, V., Meyer, S. S., … Brookes, M. J. (2018). Moving magnetoencephalography towards real-world applications with a wearable system. Nature, 555(7698), 657–661. doi:10.1038/nature26147

Brookes, M. J., Stevenson, C. M., Barnes, G. R., Hillebrand, A., Simpson, M. I., Francis, S. T., & Morris, P. G. (2007). Beamformer reconstruction of correlated sources using a modified source model. Neuroimage, 34(4), 1454–1465. doi:10.1016/j.neuroimage.2006.11.012

Dalal, S. S., Guggisberg, A. G., Edwards, E., Sekihara, K., Findlay, A. M., Canolty, R. T., … Nagarajan, S. S. (2008). Five-dimensional neuroimaging: localization of the time-frequency dynamics of cortical activity. Neuroimage, 40(4), 1686–1700. doi:10.1016/j.neuroimage.2008.01.023

Dalal, S. S., Zumer, J. M., Guggisberg, A. G., Trumpis, M., Wong, D. D., Sekihara, K., & Nagarajan, S. S. (2011). MEG/EEG source reconstruction, statistical evaluation, and visualization with NUTMEG. Comput Intell Neurosci, 2011, 758973. doi:10.1155/2011/758973

Dale, A. M., Liu, A. K., Fischl, B. R., Buckner, R. L., Belliveau, J. W., Lewine, J. D., & Halgren, E. (2000). Dynamic statistical parametric mapping: combining fMRI and MEG for high-resolution imaging of cortical activity. Neuron, 26(1), 55–67.

Dale, A. M., & Sereno, M. I. (1993). Improved localization of cortical activity by combining EEG and MEG with MRI cortical surface reconstruction: A linear approach. Journal of Cognitive Neuroscience, 5(2), 162–176.

Delorme, A., & Makeig, S. (2004). EEGLAB: an open source toolbox for analysis of single-trial EEG dynamics including independent component analysis. J Neurosci Methods, 134(1), 9–21. doi:10.1016/j.jneumeth.2003.10.009

Friston, K., Harrison, L., Daunizeau, J., Kiebel, S., Phillips, C., Trujillo-Barreto, N., … Mattout, J. (2008). Multiple sparse priors for the M/EEG inverse problem. Neuroimage, 39(3), 1104–1120. doi:10.1016/j.neuroimage.2007.09.048

Fuchs, M., Wagner, M., Kohler, T., & Wischmann, H. A. (1999). Linear and nonlinear current density reconstructions. Journal of Clinical Neurophysiology, 16(3), 267–295.

Goldenholz, D. M., Ahlfors, S. P., Hamalainen, M. S., Sharon, D., Ishitobi, M., Vaina, L. M., & Stufflebeam, S. M. (2009). Mapping the signal-to-noise-ratios of cortical sources in magnetoencephalography and electroencephalography. Hum Brain Mapp, 30(4), 1077–1086.

Gramfort, A., Luessi, M., Larson, E., Engemann, D. A., Strohmeier, D., Brodbeck, C., … Hamalainen, M. (2013). MEG and EEG data analysis with MNE-Python. Front Neurosci, 7, 267. doi:10.3389/fnins.2013.00267

Grave de Peralta-Menendez, R., & Gonzalez-Andino, S. L. (1998). A critical analysis of linear inverse solutions to the neuroelectromagnetic inverse problem. IEEE Transactions on Biomedical Engineering, 45, 440–448.

Grave de Peralta Menendez, R., Gonzalez Andino, S., Hauk, O., Spinelli, L., & Michel, C. M. (1997). A linear inverse solution with optimal resolution properties: WROP. Paper presented at the bv, Graz.

Grave de Peralta, R., Hauk, O., & Gonzalez, S. L. (2009). The neuroelectromagnetic inverse problem and the zero dipole localization error. Comput Intell Neurosci, 659247. doi:10.1155/2009/659247

Hauk, O., Stenroos, M., & Treder, M. (2019). EEG/MEG Source Estimation and Spatial Filtering: The Linear Toolkit. In S. Supek & C. Aine (Eds.), Magnetoencephalography: Springer Nature Switzerland AG.

Hauk, O., Wakeman, D. G., & Henson, R. (2011). Comparison of noise-normalized minimum norm estimates for MEG analysis using multiple resolution metrics. Neuroimage, 54(3), 1966–1974.

Henson, R. N., Mouchlianitis, E., & Friston, K. J. (2009). MEG and EEG data fusion: Simultaneous localisation of face-evoked responses. Neuroimage, 47(2), 581–589. doi:DOI 10.1016/j.neuroimage.2009.04.063

Huang, M. X., Huang, C. W., Robb, A., Angeles, A., Nichols, S. L., Baker, D. G., … Lee, R. R. (2014). MEG source imaging method using fast L1 minimum-norm and its applications to signals with brain noise and human resting-state source amplitude images. Neuroimage, 84, 585–604. doi:10.1016/j.neuroimage.2013.09.022

Iivanainen, J., Stenroos, M., & Parkkonen, L. (2017). Measuring MEG closer to the brain: Performance of on-scalp sensor arrays. Neuroimage, 147, 542–553. doi:10.1016/j.neuroimage.2016.12.048

Kietzmann, T. C., Spoerer, C. J., Sörensen, L., Cichy, R. M., Hauk, O., & Kriegeskorte, N. (2019). Recurrence required to capture the dynamic computations of the human ventral visual stream. arXiv.

Krishnaswamy, P., Obregon-Henao, G., Ahveninen, J., Khan, S., Babadi, B., Iglesias, J. E., … Purdon, P. L. (2017). Sparsity enables estimation of both subcortical and cortical activity from MEG and EEG. Proc Natl Acad Sci U S A, 114(48), E10465-E10474. doi:10.1073/pnas.1705414114

Lin, F. H., Witzel, T., Ahlfors, S. P., Stufflebeam, S. M., Belliveau, J. W., & Hamalainen, M. S. (2006). Assessing and improving the spatial accuracy in MEG source localization by depth-weighted minimum-norm estimates. Neuroimage, 31(1), 160–171.

Litvak, V., Mattout, J., Kiebel, S., Phillips, C., Henson, R., Kilner, J., … Friston, K. (2011). EEG and MEG data analysis in SPM8. Comput Intell Neurosci, 2011, 852961. doi:10.1155/2011/852961

Lütkenhöner, B. (1998a). Dipole source localization by means of maximum likelihood estimation I. Theory and simulations. Electroencephalogr Clin Neurophysiol, 106(4), 314–321.

Lütkenhöner, B. (1998b). Dipole source localization by means of maximum likelihood estimation. II. Experimental evaluation. Electroencephalogr Clin Neurophysiol, 106(4), 322–329.

Malmivuo, J. (2012). Comparison of the Properties of EEG and MEG in Detecting the Electric Activity of the Brain. Brain Topography, 25(1), 1–19. doi:DOI 10.1007/s10548-011-0202-1

Menke, W. (1989). Geophysical data analysis: Discrete inverse theory. San Diego: Academic Press, Inc.

Michel, C. M., Murray, M. M., Lantz, G., Gonzalez, S., Spinelli, L., & Grave De Peralta, R. (2004). EEG source imaging. Clinical Neurophysiology, 115(10), 2195–2222.

Molins, A., Stufflebeam, S. M., Brown, E. N., & Hamalainen, M. S. (2008). Quantification of the benefit from integrating MEG and EEG data in minimum l(2)-norm estimation. Neuroimage, 42(3), 1069–1077. doi:DOI 10.1016/j.neuroimage.2008.05.064

Oostenveld, R., Fries, P., Maris, E., & Schoffelen, J. M. (2011). FieldTrip: Open source software for advanced analysis of MEG, EEG, and invasive electrophysiological data. Comput Intell Neurosci, 2011, 156869. doi:10.1155/2011/156869

Owen, J. P., Wipf, D. P., Attias, H. T., Sekihara, K., & Nagarajan, S. S. (2012). Performance evaluation of the Champagne source reconstruction algorithm on simulated and real M/EEG data. Neuroimage, 60(1), 305–323. doi:10.1016/j.neuroimage.2011.12.027

Pascual-Marqui, R. D. (2002). Standardized low-resolution brain electromagnetic tomography (sLORETA): technical details. Methods Find Exp Clin Pharmacol, 24 Suppl D, 5–12.

Pascual-Marqui, R. D., Faber, P., Kinoshita, T., Kochi, K., Milz, P., Nishida, K., & Yoshimura, M. (2018). Comparing EEG/MEG neuroimaging methods based on localization error, false positive activity, and false positive connectivity. bioRxiv, 269753. doi:10.1101/269753

Phillips, C., Rugg, M. D., & Friston, K. J. (2002). Anatomically informed basis functions for EEG source localization: combining functional and anatomical constraints. Neuroimage, 16(3 Pt 1), 678–695.

Sarvas, J. (1987). Basic mathematical and electromagnetic concepts of the biomagnetic inverse problem. Physics in Medicine and Biology, 32(1), 11–22.

Scherg, M., & Berg, P. (1991). Use of prior knowledge in brain electromagnetic source analysis. Brain Topogr, 4(2), 143–150.

Sekihara, K., & Nagarajan, S. S. (2008). Adaptive Spatial Filters for Electromagnetic Brain Imaging. Berlin Heidelberg: Springer.

Stenroos, M., & Hauk, O. (2013). Minimum-norm cortical source estimation in layered head models is robust against skull conductivity error. Neuroimage, 81, 265–272. doi:10.1016/j.neuroimage.2013.04.086

Stokes, M. G., Wolff, M. J., & Spaak, E. (2015). Decoding Rich Spatial Information with High Temporal Resolution. Trends in Cognitive Sciences, 19(11), 636–638. doi:10.1016/j.tics.2015.08.016

Tadel, F., Baillet, S., Mosher, J. C., Pantazis, D., & Leahy, R. M. (2011). Brainstorm: a user-friendly application for MEG/EEG analysis. Comput Intell Neurosci, 2011, 879716. doi:10.1155/2011/879716

Tierney, T. M., Holmes, N., Mellor, S., Lopez, J. D., Roberts, G., Hill, R. M., … Barnes, G. R. (2019). Optically pumped magnetometers: From quantum origins to multi-channel magnetoencephalography. Neuroimage. doi:10.1016/j.neuroimage.2019.05.063

Uutela, K., Hamalainen, M., & Somersalo, E. (1999). Visualization of magnetoencephalographic data using minimum current estimates. Neuroimage, 10(2), 173–180.

Van Veen, B. D., van Drongelen, W., Yuchtman, M., & Suzuki, A. (1997). Localization of brain electrical activity via linearly constrained minimum variance spatial filtering. IEEE Transactions on Biomedical Engineering, 44(9), 867–880.

Wakeman, D. G., & Henson, R. N. (2015). A multi-subject, multi-modal human neuroimaging dataset. Sci Data, 2, 150001. doi:10.1038/sdata.2015.1

Wipf, D., & Nagarajan, S. (2009). A unified Bayesian framework for MEG/EEG source imaging. Neuroimage, 44(3), 947–966.

Woolrich, M. W., Baker, A., Luckhoo, H., Mohseni, H., Barnes, G., Brookes, M., & Rezek, I. (2013). Dynamic state allocation for MEG source reconstruction. Neuroimage, 77, 77–92. doi:10.1016/j.neuroimage.2013.03.036

Zetter, R., Iivanainen, J., Stenroos, M., & Parkkonen, L. (2018). Requirements for Coregistration Accuracy in On-Scalp MEG. Brain Topogr, 31(6), 931–948. doi:10.1007/s10548-018-0656-5

